# Dynamic self-association of archaeal tubulin-like protein CetZ1 drives *Haloferax volcanii* morphogenesis

**DOI:** 10.1101/2024.04.08.588506

**Authors:** Roshali T. de Silva, Vinaya Shinde, Hannah J. Brown, Yan Liao, Iain G. Duggin

**Affiliations:** The Australian Institute for Microbiology and Infection, University of Technology Sydney, Ultimo, NSW, 2007, Australia.

**Author notes:** Address correspondence to: Iain Duggin. School of Life Sciences, Arizona State University, USA.

## Abstract

Tubulin superfamily (TSF) proteins include the well-known eukaryotic tubulin and bacterial FtsZ families, and lesser-known archaeal CetZ family. In eukaryotes and bacteria, GTP-dependent polymerization and self-association of tubulin and FtsZ protofilaments are integral to the formation of cytoskeletal structures with essential roles in cell division, growth, and morphology. Archaeal CetZs are implicated in the control of cell shape and motility through unknown mechanisms. Here, we reveal a sequence of subcellular localization patterns of CetZ1, the prototypical member of the CetZ family, during stages of *Haloferax volcanii* rod cell development, in which it plays an essential role. Like tubulin and FtsZ, we found that CetZ1 formed GTP-dependent polymers in vitro, which appear to associate laterally as irregular polymer bundles. Mutations targeting regions predicted to mediate self-association and dynamic turnover of CetZ1, including the longitudinal (GTPase T7 and T4 loops) and lateral assembly interfaces, perturbed or altered rod shape development and subcellular assembly and dynamics, and caused corresponding effects on polymerization in vitro. Remarkably, a conspicuous amphipathic protrusion in the large microtubule (M-) loop, a characteristic of the CetZ1 subfamily, also strongly influenced function and assembly. Our findings reveal the importance of dynamic CetZ1 self-association in cellular morphogenesis involving multiple regions of the TSF fold, including tubulin- and FtsZ-like structural characteristics and CetZ1-specific features. Furthermore, they support a mechanism involving CetZ1 dynamic guidance of cell envelope-associated structures that reshape the cell during morphogenesis.

## Introduction

The cytoskeleton can serve as a scaffold and force generator in maintaining cell structure and function. Tubulin superfamily proteins are one of the main components of the cytoskeleton present in most cell types. A conserved feature of tubulin superfamily proteins is their GTP-dependent polymerization and depolymerization activity ^1,2^. Although tubulin superfamily proteins, including eukaryotic tubulin, bacterial FtsZ, and archaeal CetZ, share a common fold, their quaternary structures and associated functional roles appear to have diverged more substantially.

Eukaryotic tubulin forms microtubules which typically comprise of thirteen protofilaments formed by the polymerization of alpha and beta-tubulin heterodimers bundled into a hollow cylinder ^3^. These microtubules are essential for the formation, positioning, and dynamics of many cellular structures such as the mitotic spindle and other sub-cellular structures and organelles ^4,5^.

The prokaryotic homolog of tubulin, FtsZ, present in almost all bacteria and many archaea, does not form microtubules but instead assembles around the middle of the cell in a ring-like structure called the Z-ring to organize the cell constriction required for cell division ^6–8^. The Z ring consists of multiple FtsZ protofilaments with varying inter-protofilament distances ^9,10^ that associate with the inner surface of the cell envelope prior to the onset of cell constriction through combined activities with other division proteins ^11^.

The higher-order tubulin and FtsZ filament structures involve longitudinal (GTPase interface) and lateral self-associations that are instrumental in their function ^12^. Mutations in the longitudinal interface and GTP-binding region also strongly impact cell division. For example, the *E. coli* FtsZ.G105S mutation (also known as FtsZ84), located within the highly conserved TSF signature sequence (T4 loop), disrupts GTPase activity and Z-ring assembly in a temperature sensitive manner ^13^. Lateral associations also directly influence the stability of microtubules ^12^ and the integrity of the FtsZ-ring ^10^. For example, in *B. subtilis* FtsZ, mutations in the expected lateral association (C-terminal) domain caused extended helical filaments to form at high growth temperatures in cells, instead of normal condensed Z-rings, thus inhibiting cell division ^14,15^.

Archaea have been pivotal in understanding the activities and evolution of fundamental biological processes, particularly core information processing systems. Recently, increased focus on the diversity and cell biology of archaea has been enabled through the development of methods and experimental tools that have expanded our view of archaeal cells and their lifestyles ^16–19^. The functional and structural diversity of TSF proteins across the tree of life is arguably most evident in the archaea, many of which contain FtsZ for cell division, rare tubulin-family and noncanonical homologs of unknown functions, or members of a third archaea-specific family, CetZ, which show various sequence and structural characteristics in common with FtsZ or tubulin ^18,20^.

CetZs are distributed abundantly in haloarchaea but are also present in other groups of the Euryarchaeota, including Methanomicrobia and Thermococci ^21^. The most conserved group of CetZs across haloarchaea is CetZ1, which, in the model archaeon *Haloferax volcanii*, is required for the transition from plate to rod shaped cells. This type of morphogenic transformation occurs in a variety of conditions, including for motility, early-log phase growth, and in response to micronutrient depletion ^18,22^. CetZ1 is also expected to be essential in other conditions that promote rod development or cell elongation ^23,24^. Based on monomer and protofilament crystal structures ^18^, and the common properties of TSF proteins ^25^, CetZs are predicted to polymerize through binding GTP, forming protofilaments and potentially higher-order assemblies like those of tubulin and FtsZ. However, the function and mechanisms of CetZ polymerization and higher-order structures have not yet been identified or investigated.

Here, we began to address these gaps and used site-directed mutagenesis to identify important regions of the CetZ1 protein by assessing how the mutants impact CetZ1 function and dynamic subcellular localization, as well as polymerization characteristics in vitro. The results provided the first insights into the structure, function, and mechanisms of CetZ polymerization and support a proposed mechanism for CetZ1 involving dynamic guidance of the synthesis or assembly of other structural components of *H. volcanii* to reshape cells during rod cell development.

## Methods

### Haloferax volcanii strains and growth conditions

*H. volcanii* was grown and cultured as previously described ^22^. Briefly, all strains were grown at 42 °C using Hv-Cab or Hv-YPCab medium or on agar supplemented with uracil (50 μg/mL) to fulfil auxotrophic requirements if necessary. L-Tryptophan (2 mM) was also added to the medium to induce expression genes under the control of the *p.tna* promoter on pTA962 or pHVID9-based vectors ^19,26^. *H. volcanii* strains H26 (ID621) ^27^ and ID181 (H26 Δ*cetZ1*) ^19^ were transformed with plasmids as previously described ^28^.

### Plasmids, molecular cloning, and mutagenesis

PCR was used to generate CetZ1 point mutations. In most cases, the *cetZ1* open reading frame (ORF) was amplified in two parts, by combining end primers (Table 1, EndP) with internal primers incorporating the mutations (Table 1, as indicated). This created two fragments overlapping in the region of the mutation(s) that were joined by overlap extension PCR to create the full-length mutant ORF, before cloning. For the C-terminal tail mutant, the full-length *cetZ1* ORF was amplified in one step using a reverse primer incorporating the mutations (Table 1, CetZ1_C-term tailM_R).

To construct plasmids for *H. volcanii* expression of untagged CetZ1 point mutants, the *cetZ1* PCR fragments were gel purified and cloned into pTA962 ^26^ between NdeI and BamHI restriction sites using standard ligation and transformation procedures. The same DNA fragments were also cloned into the *E. coli* T7 expression vector pHis17 between NdeI and BamHI to make plasmids for production and purification of CetZ1 protein from *E. coli*, as described further below.

To generate plasmids for expression of CetZ1-mTurquoise2 fusions in *H. volcanii*, CetZ1 ORFs were PCR amplified from their respective pTA962-based vectors using the EndP_CetZ1_F and EndP_CetZ1_2_R primers, thus removing the stop codon. The fragments were then cloned between NdeI and BamHI sites of pHVID9 or pIDHV21 for in-frame fusion to mTurquoise2 with a flexible or semi-rigid linker, respectively ^19^. All cloned inserts were verified by Sanger sequencing and those constructed for use in *H. volcanii* were demethylated by passage through *E. coli* strain C2925 before transformation of *H. volcanii*. Lists of primers and plasmids are given in Table 1 and Table 2.

### Light microscopy

To prepare cells for imaging, cultures were grown in Hv-Cab or Hv-YPCab medium supplemented with the indicated concentrations of L-Tryptophan (Sigma) to induce expression of the cloned genes from plasmids. Cells were sampled for imaging in the early to mid-log phase (OD_600 nm_ = 0.1-0.5). A 2 μL droplet of culture was placed onto a 1 % (w/v) agarose pad containing 18 % (w/v) buffered salt water (BSW) and a #1.5 glass coverslip placed on top. Phase contrast and epifluorescence images to detect mTurquoise2 fluorescence were acquired on a DeltaVision Elite V3 inverted fluorescence microscope (GE Healthcare) with a CFP filter set (excitation = 400-453 nm; emission = 463-487 nm) and 100X 1.4 NA Plan Apo objective. Exposure and acquisition settings were maintained between strains and replicates in each experiment. Three-dimensional structured-illumination microscopy (3D-SIM) was done using a DeltaVision OMX SR (GE Healthcare) microscope with a 60X plan Plan 1.42 NA lens. Acquisition parameters were as described ^18^, using a DAPI filter for excitation (405 nm), and AF488 filter for emission (504-552 nm) for imaging of mTurquoise2. A PCO sCMOS camera was used to capture optically sectioned samples using a 125 nm z-step size.

For long-term time-lapse imaging (Figure 1d, Supplementary Video 1), the CellASIC ONIX B04A-03 Microfluidic Plate with a CellASIC ONIX microfluidic imaging system was used as described ^22^. Flow chambers were perfused at 2 psi for 20 hours with Hv-YPCab medium, the cells were imaged at 10 min intervals for 18 h at 42 °C using a Nikon Ti-E microscope with a 100X Plan Apo VC objective, a CFPHQ filter for excitation (420-445 nm) and emission (460-510 nm), and Nikon DS-Qi2 megapixel monochrome CCD camera.

**Figure 1.**
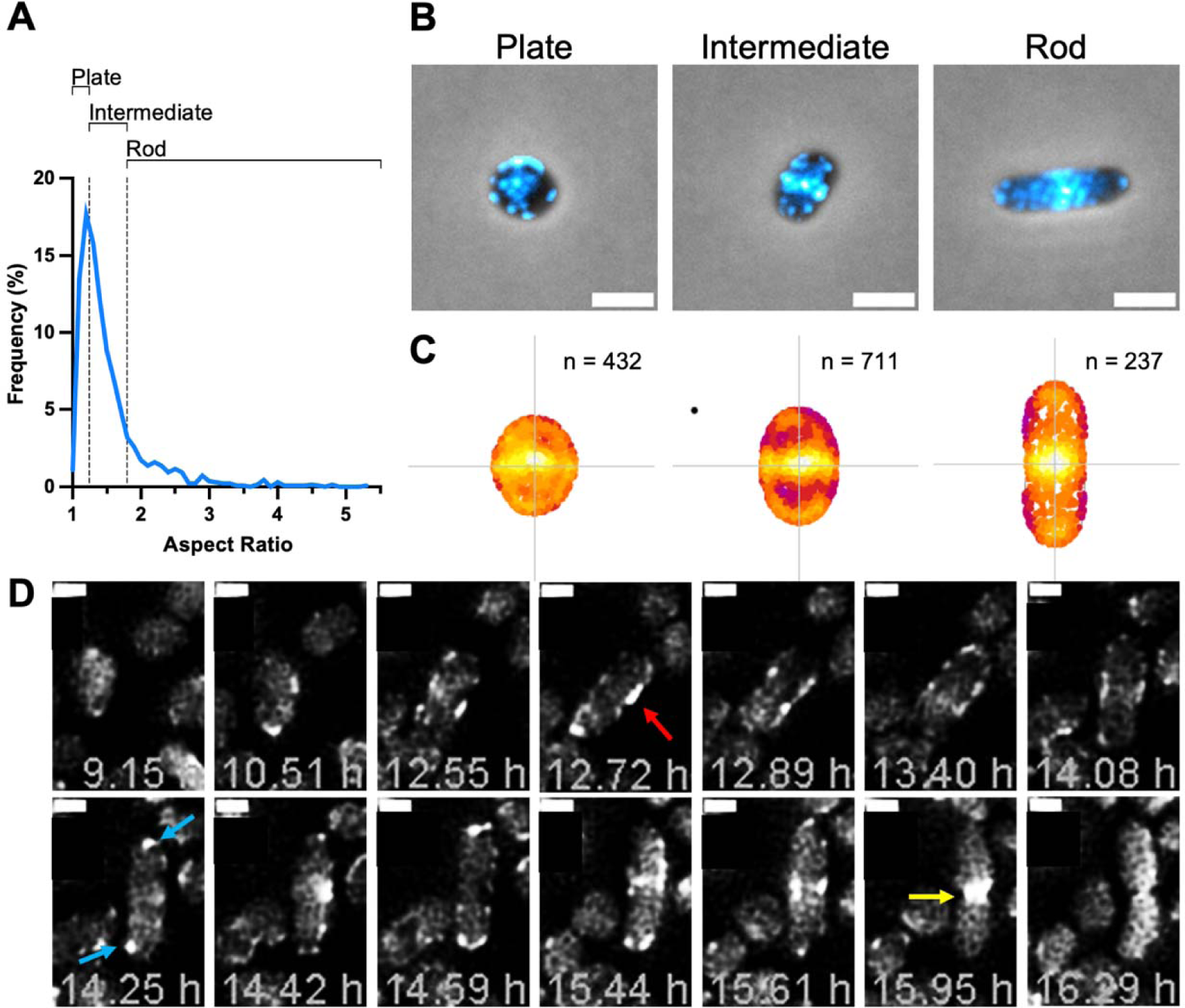
CetZ1 localization in plate, intermediate, and rod-shaped cells. *H. volcanii* ID181 (Δ*cetZ1*) background, expressing CetZ1-G-mTq2 from a plasmid was grown in Hv-Cab medium supplemented with 2 mM L-Tryptophan and sampled in early-log phase growth (OD_600_ ≈ 0.2-0.4). Data shown represent cells pooled from four culture replicates (n=1380 cells). **A)** Frequency distribution of cell aspect ratio. **B)** Representative images of CetZ1-G-mTq localization in plate (aspect ratio <1.25), intermediate (aspect ratio ≥1.25 and <1.8), and rod-shaped (aspect ratio >1.8) cells. Scale bar is 2 μm. **C)** Heatmaps of CetZ1-G-mTq localization foci in plate, intermediate, and rods. **D)** Selected timepoints throughout long-term time-lapse imaging (18 h, 10 min intervals) of CetZ1-G-mTq localization in cells transitioning from plate to rod shape. Red arrow indicates CetZ1 filaments along the long axis of the developing rod-shaped cell, blue arrows indicated polar localization of CetZ1, and yellow arrow indicates localisation of CetZ1 at mid-cell during cell division. Scale bar is 1 μm.

### Image analysis

Image analysis was done with the MicrobeJ plugin ^29^ in FIJI ^30^. Cells were identified using the default thresholding algorithm, and fluorescent foci were detected using the *point* (Figure 1c) or *foci* (Figures 4 and 5) detection methods. For both detection methods, default settings were applied except a threshold of 400 was used, and for *foci* detection a Z-score of 50 was used. Foci localisation heat maps (Figure 1c) were generated using the *XYCellDensity* function, and values for total focus intensity (i.e., the sum of pixel values above background for a region of localization defined by one maxima (focus)), the number of foci per cell, focus distance from the cell edge (Figure 4), and focus lifespan (Figure 5) were extracted from the output table. For foci tracking of time-lapse data to assess protein subcellular dynamics, raw time-lapse data were processed by registering successive images using *StackReg,* then bleach corrected (using the *Histogram Matching* algorithm) and background subtraction was applied (rolling ball size = 10 pixels) in FIJI. The limits of each focus displacement between frames were restricted to 0-0.2 μm, so that the disappearance or movement of the focus out of this region would be recorded as the end of its locational lifespan.

Raw 3-phase images generated by 3D-SIM were reconstructed using SoftWorX (Applied Precision, GE), and these reconstructed stacks were 3D rendered using the 3D script plugin ^31^ in FIJI. The *Independent transparency* algorithm was used, and all other settings were maintained as default to generate Supplementary Video 2 and 3.

### Sodium dodecyl sulphate polyacrylamide gel electrophoresis (SDS-PAGE)

For SDS-PAGE used for subsequent western blotting (Supplementary Figure 1), whole cell lysate samples were prepared by resuspension and trituration of cell pellets in a lysis buffer (20 µM Tris-HCl pH 7.4, 1 µg/mL DNaseI, 1X cOmplete™ EDTA-free Protease Inhibitor Cocktail (Roche)), followed by incubation at room temperature for 30 min. SDS sample buffer was then added to give a final (1x) buffer composition of 63 mM Tris-HCl pH 6.8, 10 % (v/v) glycerol, 5 % (v/v) 2-mercaptoethanol, 1 % (w/v) sodium dodecyl sulphate, 0.1 % (w/v) Bromophenol blue, and a OD_600_ equivalent of 5 based on the original culture sample OD_600_. Samples from protein purification and pelleting assay procedures were prepared by the direct addition of SDS sample buffer (to give a 1x final concentration) and then heated at 95 °C for 3-5 min with vortex mixing (∼1 min) to homogenize the sample. Electrophoresis was done with a 4-20% Mini PROTEAN TGX pre-cast polyacrylamide gel system (Bio-Rad) at 100 V for 85 min, including a Blue Pre-stained Broad Range Protein Standard (New England Biolabs). Gels were either stained (with 0.5% (w/v) Coomassie Brilliant Blue R in 50% (v/v) ethanol, 10 % (v/v) acetic acid, filtered), or subjected to western blotting.

### Western Blotting

Proteins were transferred after SDS-PAGE to a nitrocellulose membrane using Trans-Blot Turbo Mini 0.2 µm Nitrocellulose Transfer Packs and electro-transfer system (Bio-Rad). Total protein was stained with 0.1 % (w/v) Ponceau S (in 5 % (v/v) acetic acid) and destained with ultrapure water before imaging. The membrane was then blocked for 2 h at room temperature in 5 % (w/v) skim milk powder dissolved in TBST buffer (50 mM Tris-HCl pH 7.4, 150 mM NaCl, 0.05% (v/v) Tween-20), and then probed with anti-CetZ1 polyclonal antisera with chemiluminescence detection as described ^18^.

### Protein purification

T7 expression plasmids (pHis17-based), encoding untagged CetZ1 or its mutants, were used to transform *E. coli* C41 ^32^ via electroporation. Midlog cultures (OD_600_ = 0.4) in 2TY medium were then induced with 1 mM IPTG and incubated at 18 °C overnight with shaking. The bacteria were then harvested by centrifugation, and the cell pellet was resuspended in a buffer containing 25 mM Tris-Cl (pH 8.5), 1 mM EDTA, and 20 % (v/v) glycerol. [20 % glycerol was used to avoid CetZ1 soluble aggregate formation (non-nucleotide dependent) ^33^, which otherwise complicated purification.] The cells were lysed by sonication with sample cooling on ice. The lysate was centrifuged at 18,000 rpm for 30 min at 4 °C (Hitachi Himac R18A rotor), and then 0.2 % (w/v) polyethyleneimine was then added to the isolated supernatant. After a subsequent centrifugation at 10,000 rpm for 15 min at 4 °C, the resulting supernatant was applied to a HiTrap Q DEAE column (5 mL bed volume) running with a 0.2 μm filtered buffer containing 25 mM Tris-Cl (pH 8.5), 1 mM EDTA, and 20 % (v/v) glycerol on an AKTA Pure chromatography system (Cytiva) at 4 °C, with a gradient elution of 0-500 mM KCl. CetZ1-containing fractions, based on the UV absorbance trace (280 nm) and SDS-PAGE, were pooled, and further purified using a Superdex 200 Increase 10/300 GL column, with 25 mM Tris-Cl buffer (pH 7.5), 1 mM EDTA, 20 % (v/v) glycerol, and 200 mM KCl. Fractions containing high concentrations of CetZ1 were, pooled and concentrated using ultrafiltration and then stored at –80 °C.

### Right-angle Light Scattering Assay

A 200 µL reaction mixture was prepared containing CetZ1 or mutant protein (12 µM, unless otherwise indicated) in CetZ1 polymerization buffer containing 800 mM 1,4-Piperazinediethanesulfonic acid (PIPES) (pH 7.5), 3 M KCl, and 10 mM MgCl_2_ (prepared fresh from 0.2 μm filtered stock solutions). [The relatively high PIPES concentration was selected based on our preliminary work indicating that 0.1 M HEPES/MES/PIPES buffers (with 3 M KCl, 10 mM MgCl_2_, and 1 mM GTP) all failed to induce detected polymerization, as well as previous observations of tubulin polymerization conditions ^34^.] The mixture was then thoroughly mixed and incubated at 37 °C for 2 min ^35^. A 196 µL volume was transferred to a quartz cuvette (10 mm path length) at room temperature. This sample was used to zero a Shimadzu RF-6000 Spectrofluorophotometer set with excitation and emission wavelengths of 350 nm (slit width of 1.5 nm). Data acquisition was then commenced shortly before thoroughly mixing in 4 µL of 100 mM GTP/GDP (i.e., 2 mM final concentration). Data were recorded at 10 sec intervals for 2000 sec or until apparent signal stabilization ^35^.

### Transmission Electron Microscopy

A 50 µL sample of 12 µM CetZ1 or mutants in CetZ1 polymerization buffer was prepared and then GTP or GDP was added (to 2 mM) and mixed and incubated for various times up to 35 min at 30 °C with 300 rpm shaking. A 2 µL volume was then applied to a glow-discharged formvar-carbon film coated copper EM grid (400-mesh, ProSciTech) and incubated at room temperature for 1 min. The liquid was removed by touching the edge to filter paper, and then 4 µL of a 2 % (w/v) uranyl acetate solution (0.2 μm filtered) was added. Excess stain was removed using filter paper, and the grid was allowed to dry before imaging with a Tecnai T20 microscope operating at 120 kV. Images were captured with a Gatan CCD camera.

## Results

### CetZ1 displays distinct subcellular localization patterns throughout rod cell development

CetZ1 tagged with GFP displays a complex localization pattern in *H. volcanii*, with foci, patches and filaments appearing around the cell envelope often near the cell poles and mid-cell regions ^18^. In recent work, we generated a new C-terminal fusion between CetZ1 and the mTurquoise2 (mTq2) fluorescent protein, which supported a high level of function in cell motility ^19^. In the current study, we started by observing CetZ1-mTq2 subcellular localization patterns from samples taken during early-log phase growth when many cells are transitioning to rod-shape. Based on the distribution of their aspect ratios, we divided cells into three categories (Figure 1a): plates (aspect ratio <1.25), intermediates (aspect ratio ≥1.25 and <1.8), and rods (aspect ratio ≥1.8). CetZ1 localization differed in all three shape categories (Figure 1b, c). In plates, CetZ1 localized in small patches primarily around the cell outline distributed throughout the cell. In intermediates, CetZ1 localization became more concentrated around the cell outline and at mid-cell. In rod-shaped cells, mid-cell and polar localization became more prominent. Since these differing patterns appeared to correlate with cell aspect ratio, they likely reflect stages or distinct activities required for rod-development.

To further investigate, we conducted time-lapse imaging of cells expressing CetZ1-mTq2 as they transition from plates to rod-shaped cells in an incubated microfluidic chamber system (Figure 1d, Supplementary Video 1). Although the CetZ1-mTq2 localization dynamics were too rapid to be motion-tracked at the frame rate selected to minimize photobleaching over the rod development time course (10 min intervals over 18 h), the results revealed a sequence of patterns of CetZ1-mTq2 localization that typically appeared during rod-development. We observed that plate shaped cells, prior to obvious rod formation, displayed a patchy localization much like the CetZ1-GFP patterns seen previously in plates sampled during mid-log growth (Figure 1b) ^18^. As the cells began to elongate and narrow, CetZ1 often appeared as short filaments along the edges of the cell’s long axis (Figure 1d, red arrow), and then towards the cell poles as patches or end-cap structures in the rods (Figure 1d, cyan arrow). Subsequently, before and during rod cell division, CetZ1 was often seen concentrated around mid-cell, particularly during constriction, but was typically more distributed around a broader midcell region (Figure 1d, yellow arrow) compared to the condensed constricting rings seen with *H. volcanii* FtsZ1 and FtsZ2 ^8^.

### Probing the role of CetZ1 domains in self-association

The sequence of dynamic patterns of CetZ1 localization (Figure 1d) suggested that CetZ1 likely forms different assemblies during a staged rod development process, in which it is known to play and essential role ^18^. Since very little is known about CetZ family higher-order assemblies and their dynamics, which currently are largely inferred from knowledge of other TSF proteins, this prompted us to begin investigating CetZ1 polymerization at the molecular level with the goal of defining regions involved in polymerization or its regulation and their roles in cellular function and localization. We designed site-directed mutants of CetZ1 in regions of predicted self-association, or other potential interactions, based on amino-acid and domain conservation amongst CetZs and well-characterized TSF members, with close reference to existing crystal structures of the *H. volcanii* CetZ1 monomer and a CetZ2 polymer-like form ^18^. The CetZ2 multimer shows a 2D sheet-like structure with longitudinal and lateral self-associations that resemble the arrangement of tubulin within the microtubule wall as well as FtsZ protofilaments and their expected lateral association ^12^.

Alphafold Multimer was used to generate a predicted polymer structure containing nine monomers of CetZ1, displaying the positions of the mutations (Figure 2). As expected, the structure strongly resembled the experimental structures of CetZs and other tubulin superfamily proteins, with both longitudinal and lateral self-associations. It also includes surface regions that were not resolved in the CetZ crystal structures but were targeted as part of our set of mutants designed to probe potential interaction interfaces due to their conservation, chemical properties, or location, as described below.

**Figure 2.**
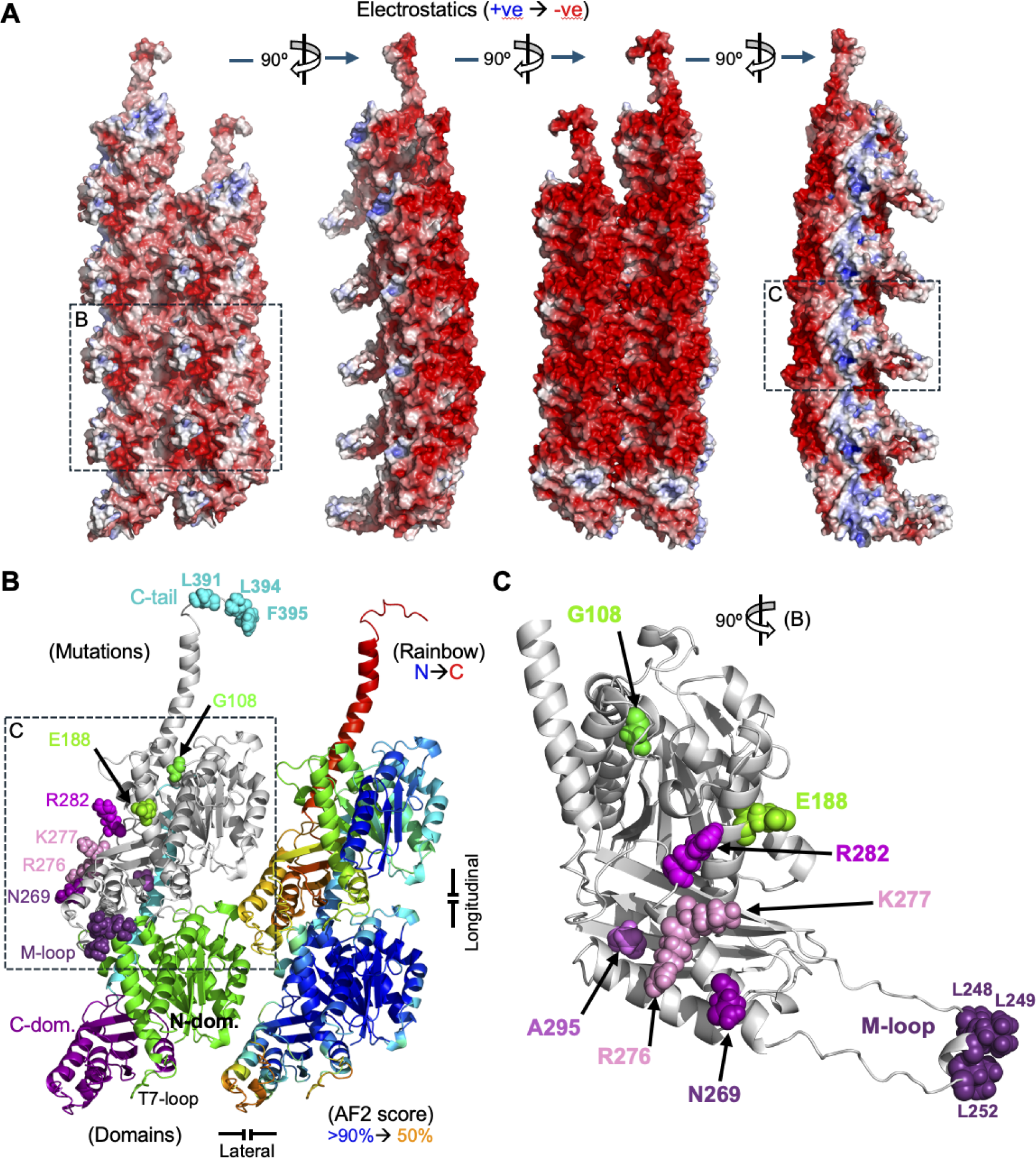
Model of CetZ1 sheet-like polymers indicating residues mutated. Nine monomers of *H. volcanii* CetZ1 were input for Alphafold2 Multimer structure prediction. **A)** The most probable model shown with surface electrostatics has two laterally associating protofilaments. Grey indicates hydrophobic surfaces. Weaker predictions showed single protofilaments arranged in rings (not shown), suggesting the potential for significant flexibility in protofilament conformation. **B)** Four CetZ1 monomers in the predicted polymer, each shown with a different colour scheme indicated in parentheses. Residues mutated in this study are shown as space-filling models (N-terminal domain – greens, C-terminal domain – purples, and C-terminal tail – cyan). The C-terminal tail and M-loop mutations had 3 amino-acid substitutions each, to reduce hydrophobicity from these regions (see text). The longitudinal and lateral subunit repeat distances are 4.2 nm and 4.5 nm, respectively. The GTPase active site is generated by both subunits at of the longitudinal interface. **C)** CetZ1 monomer from the filament, with the predicted lateral association regions facing the viewer.

In the tubulin superfamily, the longitudinal interface is largely responsible for binding and hydrolysis of GTP to control polymer stability ^25^. In the CetZ1 N-terminal GTP-binding domain (Figure 2b, green) we made a mutation (G108S) in the ‘signature sequence’ of the tubulin superfamily (in the T4 loop of the GTP-binding pocket) analogous to an *E. coli* FtsZ mutation (G105S, or FtsZ84) that impairs GTPase functions and polymer assembly and stability ^13^. CetZ1.G108S is expected to produce differing effects on polymerization and function compared to the expected GTPase-blocked E218A (T7 loop) mutant ^18^, which is further investigated here. We also created the E188G mutation on the ‘front’ surface of the predicted CetZ1 sheet in Figure 2b (comparable to the inner surface of a microtubule), which is not in direct contact with other subunits or the GTP-binding pocket in the model but might interfere with potential interactions of the front surface with other molecules in vivo.

In the CetZ1 C-terminal domain, expected to participate in polymerization and lateral association, we created three site-directed mutants to probe the external surface of this domain (N269A, R282A, and R276E/K277E), where a noticeable region of partial positive charge exists in the otherwise highly negatively charged CetZ1 protein in the region of the predicted lateral associations (Figure 2). We also created a mutation (A295S) within the C-terminal domain, based on a region of FtsZ mutations in *B. subtilis* that impair lateral association and Z-ring condensation at high growth temperatures ^14^.

A distinctive characteristic of the CetZ1 subfamily is an unusually long loop in the C-terminal domain that conspicuously protrudes from the surface and has a hydrophobic cluster of amino acids at the tip that might be part of a short amphipathic helix (Figure 2b). This corresponds to the shorter microtubule (M-) loop in tubulin, which influences lateral associations between protofilaments ^12^. We therefore sought to investigate the importance of the conspicuous hydrophobicity in the CetZ1 M-loop, which might form an amphipathic helix (Figure 2), by substituting three leucine residues (LLSRL to SASRA) (Figure 2c). Lastly, the very C-terminal amino acids of CetZ1, as with other tubulin superfamily proteins, contains a short, conserved sequence implicated in mediating protein-protein interactions, which in CetZ1 shows exposed hydrophobic residues. The C-terminal tail also appears to associate with the adjacent subunit, on the ‘back’ or outer surface with respect to microtubule structure, and might therefore be involved in polymerization (Figure 2) ^18^. We made substitutions at the very C-terminus of CetZ1 (LESLF to AESGG) to begin investigating its potential role.

### CetZ1 site-directed mutants have differing effects on cell shape

To begin to determine the effects of the site-directed mutations on CetZ1 function, the mutants and a wild-type control were expressed from respective plasmids in a *cetZ1* in-frame knockout background (ID181) and initially screened for their capacity to support rod cell formation. Cells were grown in rich medium (Hv-YPCab) supplemented with 2 mM L-Tryptophan (Trp) to induce expression of the wild-type or mutant genes, and samples were taken in the early-log phase for imaging (Figure 3). Circularity analysis of individual cells was used to quantify the effects of each mutation on cell shape (Figure 3b). Under these conditions, wild-type *H. volcanii* appears pleomorphic and a substantial fraction of cells are rod shaped, reflected in a reduced circularity. As expected, rods were absent in the *cetZ1* knockout (ID181 + pTA962 vector only) but expression of wild-type *cetZ1* from the plasmid partially complemented the rod defect (Figure 3b).

**Figure 3.**
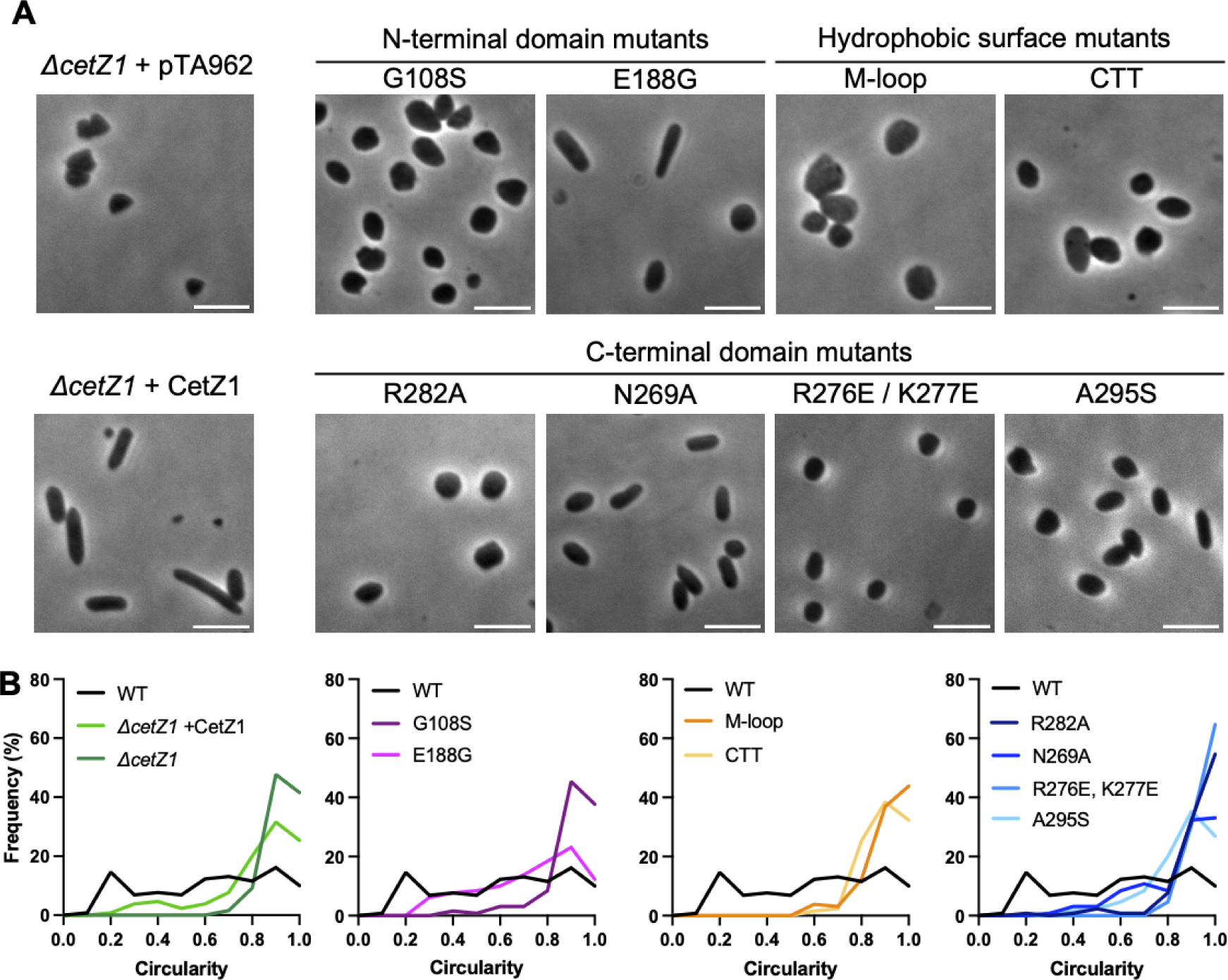
Effects of CetZ1 point mutations on cell shape. CetZ1 mutants were expressed in *H. volcanii* ID181 (Δ*cetZ1*) and cells were grown in Hv-YPCab medium supplemented with 2 mM L-Trp, before sampling in early log phase for phase-contrast microscopy. **A)** Representative phase contrast images of the relevant *H. volcanii* wild-type (WT, H26+pTA962) and (ID181+pTA962) control strains (pTA962 – vector only), and ID181 expressing the indicated *cetZ1* point mutants. Scale bar = 5 μm. **B)** Cell circularity analyses of individual cells shown as frequency distributions (1 = circular outline). The same data for WT is included in all plots for reference. 130 individual cells were analysed for each strain.

All but one of the mutants had an impact on rod cell development. The N-terminal domain mutant, G108S, failed to support rod formation; almost all cells showed a circularity > 0.8, representing the discoid or ‘plate’ shaped cell type, like the *cetZ1* knockout. Whereas the E188G mutation, located outside the predicted longitudinal and lateral interfaces, supported rod development like the wild type (Figure 3b). On the other hand, the surface hydrophobic patch mutations in the M-loop and the C-terminal tail (CTT) that are also outside the primary longitudinal and lateral interfaces, failed to support rod development (i.e., cells with circularity < ∼0.7), indicating that these regions are critical to CetZ1 function. The C-terminal domain mutations at the predicted lateral interface had varied effects; R282A and R267E/K277E that contribute to the positively charged region in the center of the predicted lateral interface (Figure 2) strongly blocked rod development, whereas N269A and A295S in the vicinity had partial effects (Figure 3b).

### CetZ site-directed mutants have distinct effects on subcellular localization

Given that most of the mutants we made to the surface of the core fold of CetZ1 showed a strong impact on rod cell development, we chose to further investigate the role of the CetZ1 GTP binding (N-terminal) and predicted (C-terminal) domains in cellular localization and polymerization functions, which had not yet been investigated in the CetZ family but are integral to those functions in other TSF proteins. We tagged the selected CetZ1 mutants with mTq2 to observe their localization patterns in early-log phase growth (Figure 4a) and used western blotting to check the production and integrity of each protein (Supplementary Figure 1). We then quantitatively analyzed the localization patterns (Figure 4b-d).

**Figure 4.**
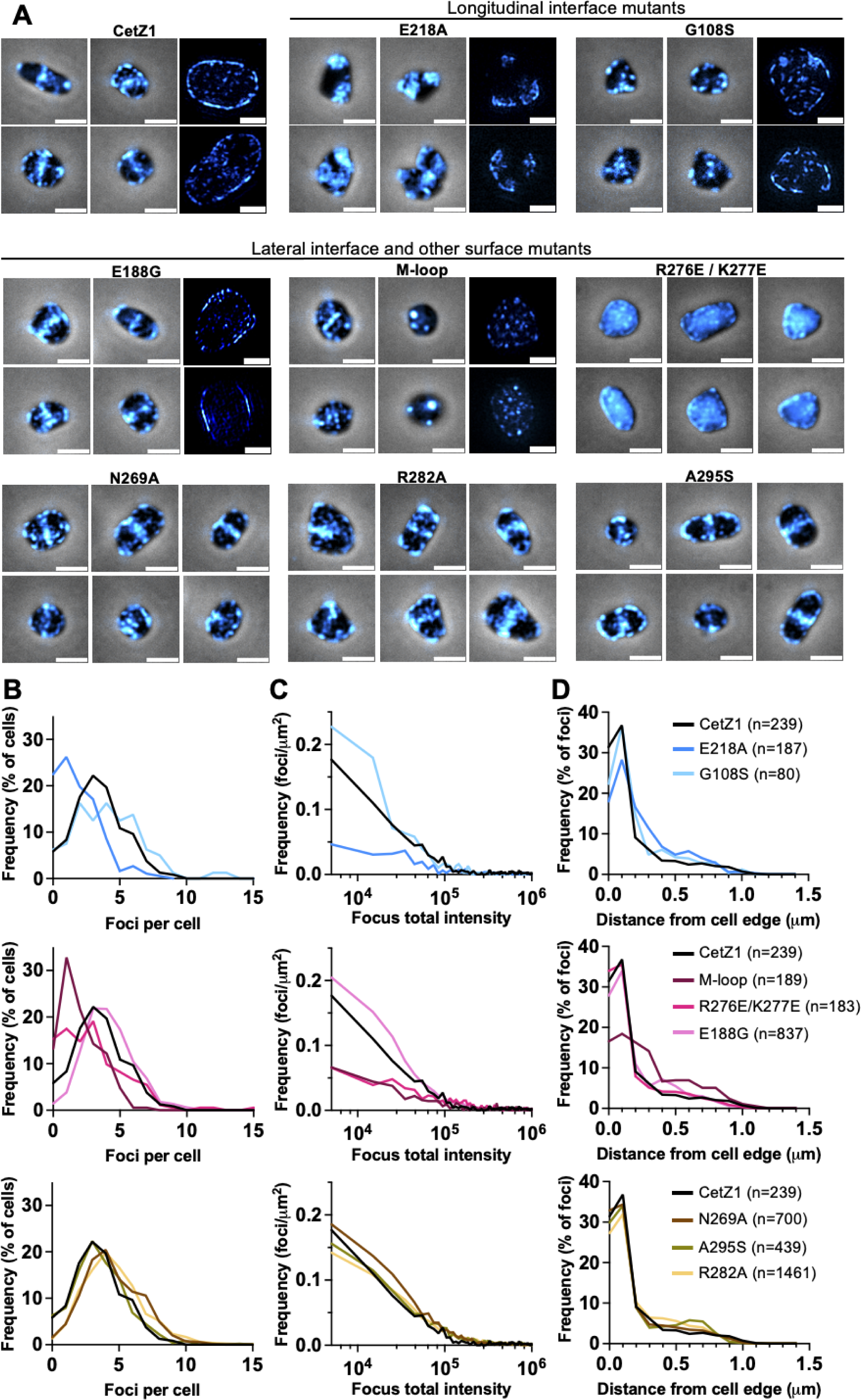
Point mutations in CetZ1 affect subcellular localization patterns. CetZ1 wild-type or the indicated mutants were expressed in the *H. volcanii* ID181 (H26 Δ*cetZ1*) background by growth in Hv-Cab medium supplemented with 2 mM Trp before imaging in the early log phase. **A)** Phase-contrast and epifluorescence microscopy composites of CetZ1 mutants tagged with mTq2 (scale bar = 2 μm), and 3D-SIM (righthand two images for CetZ1, E218A, G108S, M-loop, and E188G - scale bar = 1 μm). Exposure settings were maintained between all strains. All images shown were subjected to background subtraction in FIJI using a rolling-ball size of 10 pixels except R267E/K277E due to its obvious diffuse distribution. Frequency distributions are shown of **(B)** the number of localizations per cell (‘foci’ here refers to localizations of any shape), **(C)** the focus total intensity, and **(D)** the distance of each focus center to the closest part of the cell outline. Data for wild-type CetZ1 is included in all plots for reference. The key and number of cells (n) applies to all panels.

Amongst the three N-terminal domain mutants, G108S, E188G, and E218A, previous work on E218A (in the T7 loop) showed that it inhibits rod cell development and generates a ‘jagged’ cell phenotype in a wild-type *H. volcanii* background, characterized by short cell protrusions and regions of high local cell envelope curvature, where it tends to localize ^18^. Here, with the mTq2 tag and ID181 (Δ*cetZ1*) background, we observed consistent results; E218A typically appeared as relatively fewer larger and brighter patches, especially at regions of greater curvature compared to the wild-type protein (Figure 4a). In comparison, we saw that the other mutation in the GTP-binding interface, G108S (in the T4 loop), appeared to produce many smaller foci and short patches or filaments as well as some brighter localizations (Figure 4a). Super-resolution 3D-SIM imaging showed these in greater detail and suggested that the E218A and G108S GTP-binding domain mutants are associated with the cell outline to approximately a similar degree as the wild-type CetZ1 (Figure 4a, top righthand sections). Quantitative image analysis of the epifluorescence data (Figure 4b, 4c top) verified that E218A had fewer localizations (foci) compared to the wild-type, consistent with its expected effect of hyper-stabilization of polymers ^18^, whereas G108S showed more of the dimmer foci than the wild-type (Figure 4c, top) consistent with potential destabilization (fragmentation) of G108S polymers. Analysis of foci proximity to cell outline, showed these two mutants had a similar or slightly greater foci-to-outline distance compared to wild-type CetZ (Figure 4d).

The fully functional mutant, E188G, on the ‘front’ surface of the N-terminal domain (Figure 2b), appeared like the wild-type when observed by epifluorescence and 3D-SIM (Figure 4a), with minor differences in the distribution of foci frequency and intensity (Figure 4a and 4c, middle). Cell outline association of E188G was indistinguishable from the wild type (Figure 4d, middle). In stark contrast, the M-loop mutation that protrudes from the front surface of the C-terminal domain had a striking effect on localization (Figure 4a), with only one or a few bright isolated circular foci per cell, with no clear evidence of filaments or patch-like structures, and cellular background fluorescence that varied between samples and individual cells (Figure 4a-c). Interestingly, the M-loop mutant foci were strongly disassociated from the cell outline compared to wild-type and other mutants (Figure 4a, 4d - middle). In agreement, 3D-SIM reconstructions clearly showed the M-loop mutant foci distributed throughout the cytoplasm (Figure 4a, Supplementary Video 3).

Mutations targeting the predicted lateral association interface showed significant differences. The most substantive mutation in the centre of the predicted interface, R276E/K277E, had a strong impact on localization, with most cells displaying predominantly diffuse signal and relatively dim foci (Figure 4b-d, middle), consistent with a major defect in polymerization but without the aggregation apparent for the M-loop mutant. The other C-terminal domain mutants in the vicinity of predicted lateral interface (Figure 4a, bottom row) were not dramatically different to wild type (Figure 4a-c, bottom), although R282A showed a noticeable propensity to line regions of high curvature of the cell outline (Figure 4a, centre bottom). These results therefore indicate that the CetZ1 C-terminal domain and especially the isolated positively charged region at the direct-contact interface (Figure 2), play an important role in CetZ1 assembly in vivo.

### CetZ point mutants with altered subcellular patterns affect protein dynamics in vivo

CetZ1 and selected mutants with different localization patterns and those predicted to influence polymer turnover via GTPase activity, i.e., E218A, G108S, M-loop, and E188G, were next analyzed by time-lapse imaging over 30 min (Figure 5a, Supplementary Videos 4-8). CetZ1 exhibits complex dynamic patterns in vivo, so we quantified and compared overall dynamics by detecting fluorescent foci and measuring their locational lifespans, i.e., the time until they disappear or move more than 200 nm between the 1 min intervals. As may be seen in Figure 5b, the E218A, G108S and M-loop mutants had a greater frequency of relatively long-lived foci, whereas E188G showed slightly fewer foci with intermediate lifespans, compared to the wild-type.

**Figure 5.**
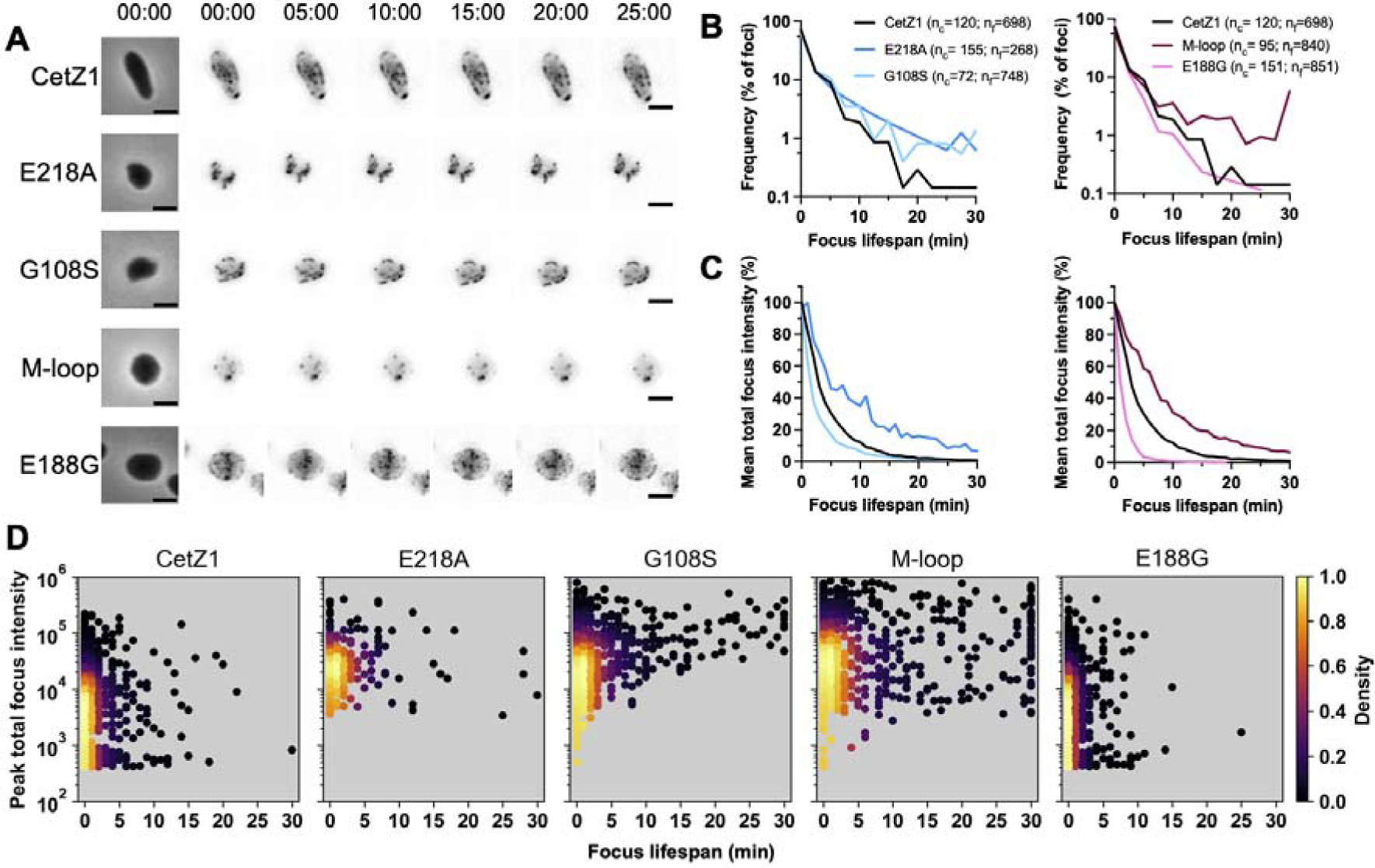
In vivo dynamics of CetZ1 and CetZ1 mutants. **A)** Time-lapse imaging (1 min intervals for 30 min) of *H. volcanii* ID181 expressing the indicated mutants tagged with mTq2, grown in Hv-Cab medium with 2 mM Trp. Scale bar = 2 μm. **B)** Frequency distributions of focus locational lifespans. Foci were recorded as ‘alive’ if detected above the threshold and within a 200 nm radius between frames. Bin size = 2. **C)** Mean of total focus intensity, over focus locational lifespan. The total intensity of each focus was normalized to 100% at its ‘birth’. The same data for CetZ1 is shown in both plots of B and C for reference. **D)** XY scatter plots showing the maximum (peak) focus total intensity reached compared to its total lifespan. Plots are colored by normalized density, reflecting the frequency of foci at each position. The total numbers of individual cells analyzed (n_c_) and detected foci within them (n_f_) shown in panel B refers to panels B-D. Note that the 30 min endpoint graphed represents foci with lifespans of 30 min or greater, and measurement of other foci lifespans will be limited by photobleaching.

The total intensity value of each focus was then normalized as a percentage of its initial value, and the mean for each strain over locational lifespan was graphed, thus reflecting the rate of disappearance of foci (Figure 5c). This showed that the G108S and E188G localizations disappeared sooner than wild-type CetZ1 on average, whereas the E218A and M-loop were more stable, independent of foci size. Since G108S increased the frequency of the most long-lived foci (Figure 5b), the combined results indicate that while some G108S foci are hyperstable, the majority are hypostable. Furthermore, as G108S showed a greater frequency of faint foci compared to wild-type (Figure 4c), they appear to be the hypostable ones.

We therefore investigated the relationship between focus maximum intensity and its lifespan (Figure 5d). The functional proteins, CetZ1 and E188G, showed similar distributions dominated by relatively low intensity foci, the majority of which showed dynamic turnover within the first few minutes of their detection. E218A, which is expected to block GTPase activity and causes intense hyperstable localizations ^18^, showed much brighter and longer-lived foci on average, as expected. Interestingly, G108S showed a correlation between focus intensity and lifespan that confirmed the observations above, revealing the existence of bright longer-lived foci, as well as faint shorter-lived foci that never existed for longer than ∼1-2 min (Figure 5d). By comparison, wild-type CetZ1 showed greater stability of the faint foci, but more dynamic turnover of the brighter ones, and no clear correlation between intensity and lifespan (Figure 5d). A broad dynamic range of both intensity and lifespan was seen with the M-loop mutant, which showed some very long-lived (> 30 min) bright foci, consistent with their appearance as unstructured aggregates postulated above (Figure 4a, 5d, Supplementary Video 3 and 7).

### Self-assembly of CetZ1 in vitro

Full-length CetZ1 (untagged) was overproduced in *E. coli* and purified, with the aim of identifying and characterising CetZ1 self-assembly, and enabling comparisons with the mutant proteins and their activities in vivo. We found that the presence of 20% glycerol avoided soluble small aggregate formation (non-nucleotide dependent) and allowed the purification of monomeric CetZ1 (Figure 6a). To determine whether CetZ1 is capable of GTP-dependent self-assembly, a 90° light scattering assay was used for real-time measurement of polymerization. We identified buffer conditions supporting GTP-dependent CetZ1 polymerization, which included a 0.8 M PIPES buffer, as observed for tubulin ^34^, and 3 M KCl that approximates the in vivo ionic conditions (Figure 6b). The same conditions but with GDP instead of GTP, or BSA instead of CetZ1, failed to cause any detectable light scattering (Figure 6c). Consistent with this, an ultracentrifugation assay showed that CetZ1 could be recovered in the pellet (polymers) in presence of GTP but not GDP (Supplementary Figure 3a).

**Figure 6.**
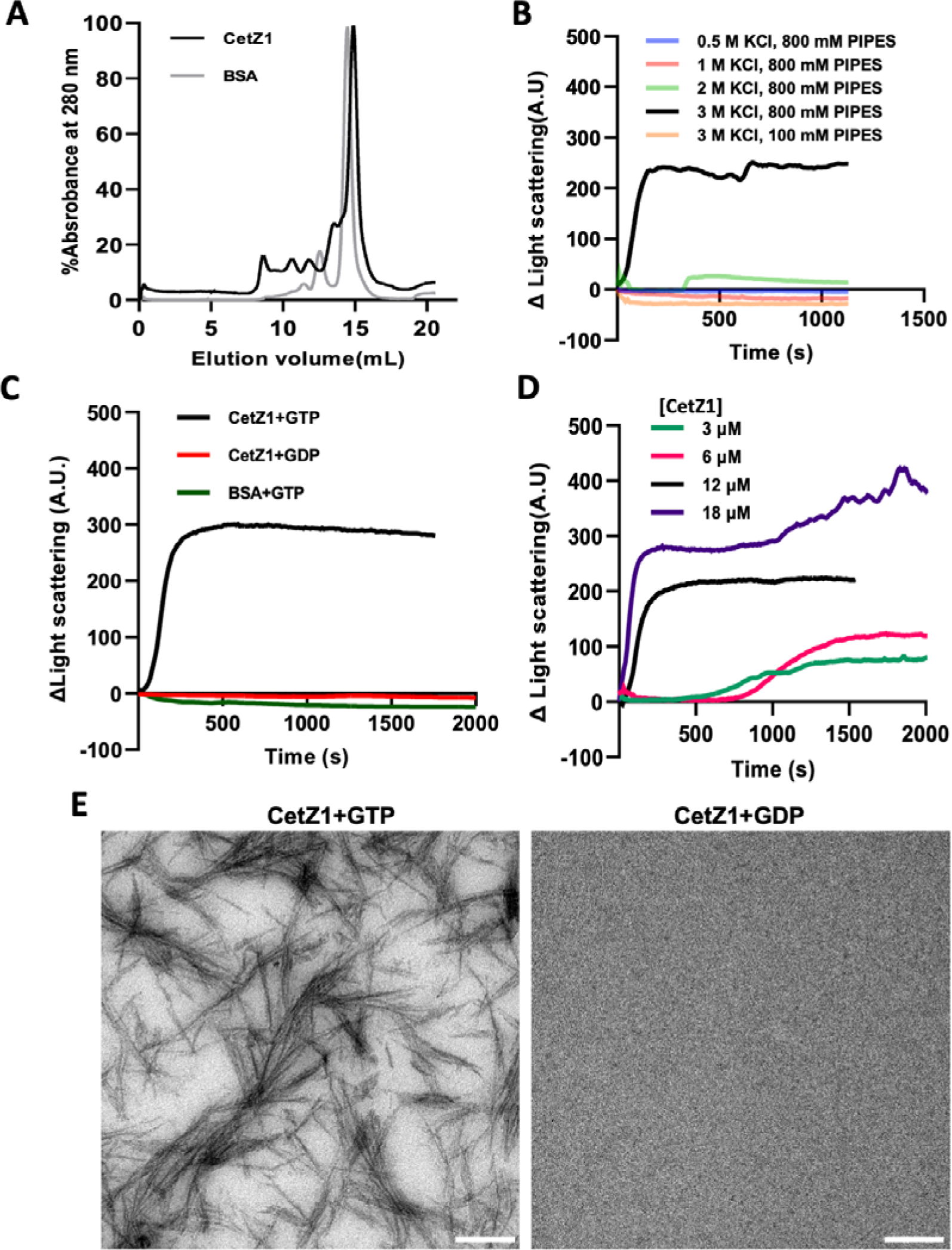
In vitro characteristics of CetZ1. **(A)** Gel filtration chromatograms of BSA (66 kDa) and final stage purification sample of CetZ1 in a buffer including 20 % (v/v) glycerol. The highest peak at ∼15 mL was isolated for polymerization experiments. **(B)** CetZ1 polymerization was examined by a 90° light scattering assay under various KCl and PIPES buffer (pH 7.5) concentrations. **(C)** As controls, GDP and BSA were added at the same concentration instead of GTP and CetZ1, respectively. **(D)** Scatter profiles with different concentrations of CetZ1. **(E)** TEM images depicting polymerized CetZ1 filaments formed in the presence of GTP after 2 minutes of incubation and non-polymerized CetZ1 in the presence of GDP (scale bar = 100 nm.)

Experiments testing the CetZ1 concentration dependence of polymerization are shown in Figure 6d. At 3 µM and 6 µM CetZ1, a lag was observed and the increase in scattering proceeded relatively slowly and to a reduced extent compared with 12 and 18 µM CetZ1, where the scattering signal rapidly increased and reached a plateau within ∼5 min. This suggests that cooperative polymerization initiates at or above concentrations approaching 12 µM CetZ1.

We then visualized the reaction samples by uranyl acetate staining and transmission electron microscopy. This revealed filaments that formed in a GTP-specific manner (Figure 6e). Some filaments were bundled together adjacently or in curved and clustered forms. The distinct filaments were ∼3 nm wide and the distance between centres of adjacent filaments was ∼4 nm, consistent with expected sizes of CetZ1 protofilaments and the CetZ2 sheet-like crystal structure ^18^. The filaments appeared to be stained directly, rather than negatively, which might be caused by the highly negative surface charge of CetZ1 (Figure 2) associating with the UO_2_^2+^ ions. Alternatively, the filaments might be uranyl phosphate (or a related salt) needle crystals formed after phosphate release by CetZ1-mediated GTP hydrolysis. Hence, we next assessed polymerization by the CetZ1 GTPase domain mutants, with the aims of clarifying filament composition and comparing their basic assembly properties.

### In vitro polymerization of CetZ1 mutants targeting the CetZ1 core fold

We purified selected mutants targeting the expected longitudinal (E218A, G108S) and lateral (M-loop, R276E/K277E) interfaces, as well as E188G. All these mutants showed GTP-dependent polymerization, with substantial differences in the rates, extent, and stability compared to wild-type CetZ1 (Figure 7a). E218A showed a relatively slow increase in scatter compared to wild-type CetZ1 but continued to increase over at least 2000 sec (Figure 7a, left), suggesting that this mutation causes a partial impediment to the rate of polymerisation, and leads to the formation of hyperstable polymers. In contrast, G108S showed limited rapid polymerization initially, followed by a gradual disassembly, suggesting that this mutation reduces the overall stability of the polymers formed in vitro. The R276E/K277E and E188G mutants exhibited reduced polymerization rates and ∼3-4 fold less peak light scattering compared to wild-type (Figure 7a, right). The M-loop mutant showed a drastic polymerization defect, with a brief pulse of scattering signal followed by a rapid and complete reversal (Figure 7a).

**Figure 7.**
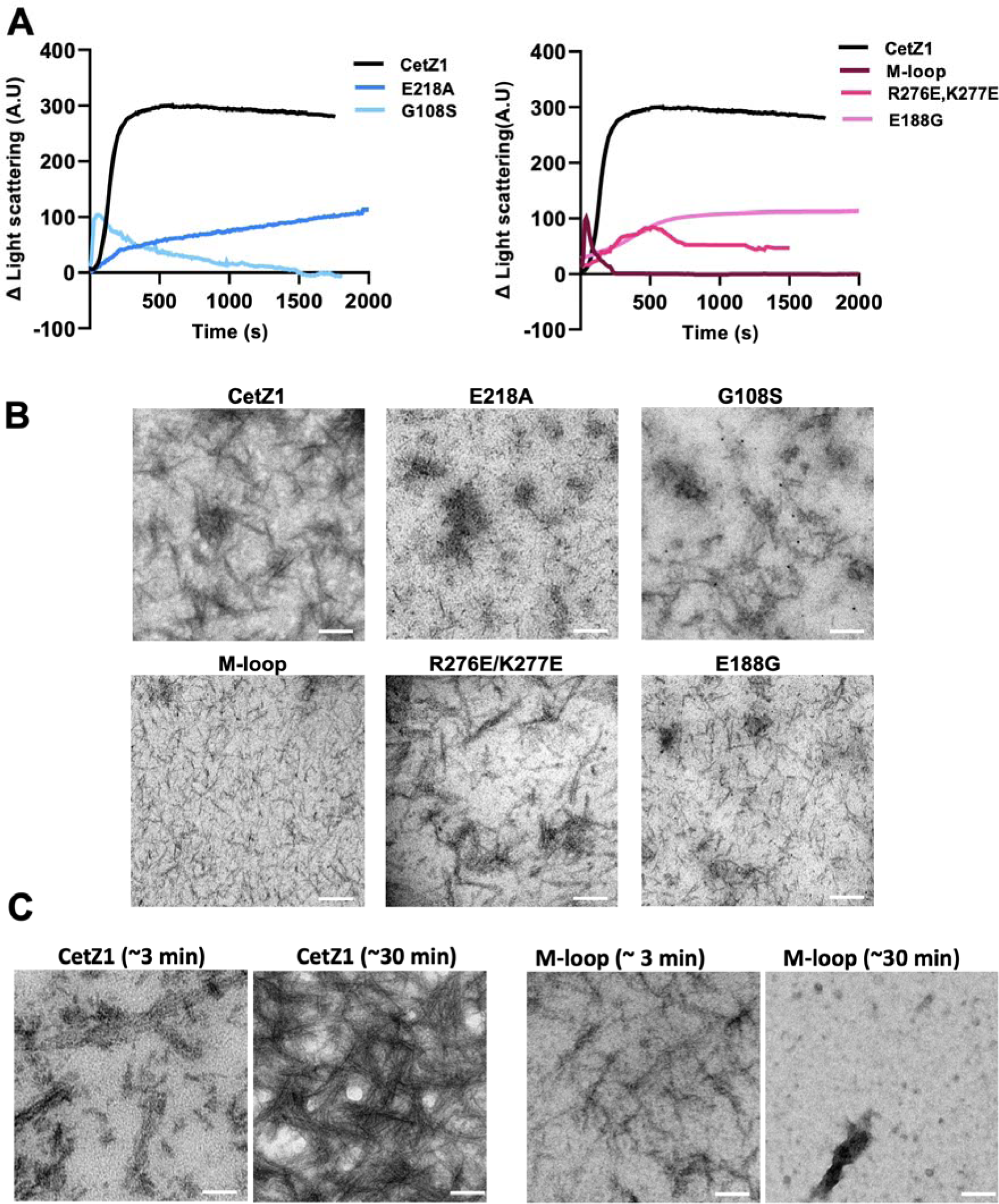
In vitro polymerization of CetZ1 core domain mutants. **(A)** Polymerization of CetZ1 mutants with right-angle light scattering. The same WT data are shown in both graphs for reference. **(B)** TEM of wild-type CetZ1 and the indicated mutant polymerization reactions with GTP after 2 min of incubation. Scale bars = 50 nm. Filament lengths were measured; bars represent the mean and standard deviation. A one-way analysis of variance (ANOVA) showed a significant difference (p<0.0001) of all mutants compared to the wild type. **(C)** TEM images of CetZ1 and M-loop mutant samples after ∼3 min and ∼30 min of adding 2 mM GTP in CetZ1 polymerization buffer. Scale bars = 50 nm. Equivalent GDP reactions with light scattering and TEM showed no evidence of polymerization with any of the mutants.

TEM of samples taken soon after the addition of GTP were generally consistent with this, where the E218A and G108S appeared as fewer, smaller filaments, E188G formed relatively short filaments consistent with its slower assembly profile (Figure 7a) and R276E/K277E gave a range of sizes, suggesting this region is not essential for detection of filaments, at least under the conditions identified here (Figure 7b). Importantly, filaments in the M-loop mutant sample were only detected soon after the addition of GTP, in both the scattering and TEM analyses (Figure 7a, 7c). This behaviour, and the differing appearances of the GTPase domain mutants, supports the interpretation that the observed filaments are protein polymers.

Given the above results, and the non-envelope associated apparent aggregation of the M-loop mutant in vivo (Figure 4), we purified *H. volcanii* lipids and investigated its potential membrane association by liposome ultracentrifugation pull-down tests. Both CetZ1 and M-loop mutant associated similarly with liposomes in a non-nucleotide dependent manner (Supplementary Figure 3b). If such membrane association is physiologically relevant, the M-loop tip hydrophobic residues mutated here therefore appear to be not essential for this membrane association, at least in vitro, and may instead be primarily involved in important lateral or longitudinal subunit interactions, analogous to the function of the M-loop in microtubules.

## Discussion

This study investigated the dynamic localization and self-association behaviour of CetZ1, a tubulin-like protein crucial for cellular morphogenesis in the archaeon *Haloferax volcanii*. Spatiotemporal imaging of functional tagged CetZ1 during rod development in vivo showed dynamic assembly behaviour and a typical sequence of characteristic patterns seen during rod development, including envelope-associated filaments along the long axis of developing rods, end-caps and polar patches in newly formed rods, and midcell localization during subsequent rod cell division. This implied that CetZ1 function involves dynamic repositioning and repurposing during stages of rod cell development.

To begin characterizing the molecular basis for CetZ1 function and the role of the dynamic structures, we identified conditions for GTP-dependent polymerisation of CetZ1 in vitro, which included a 0.8 M PIPES buffer, similar to tubulin ^34^, and a requirement of 3 M KCl, consistent with the *in vivo* ionic environment of *H. volcanii* and other investigations of halophilic protein function ^36,37^. CetZ1 also showed evidence of cooperative polymerization, as seen in other TSF proteins ^38^, and appeared to form straight or slightly curved protofilaments and irregular bundles by TEM. We then examined the impact of site-directed mutants on function and polymerization, which were designed to disrupt predicted longitudinal and lateral self-associations and other CetZ1-specific features.

Previous work had suggested that GTPase-dependent polymerization would be important in CetZ1 function, because mutation of a known catalytic residue of the T7 loop in TSF proteins (E218A), blocked cell motility and rod development and stabilized subcellular localization ^18^. Here, we expanded on this with in vitro studies of protein polymerization and by applying CetZ1 fluorescent fusions with improved function as the sole copy in vivo. We also introduced another mutation in the N-terminal GTP-binding domain, G108S (in the T4 loop near the guanine base), which is homologous to the substitution in *E. coli ftsZ84* (G105S) that affects GTPase function and longitudinal interactions ^13,39^.

The CetZ1 E218A mutation caused an impediment to polymerization in vitro but also hyperstabilized its assembled structures in vivo and in vitro, consistent with its expected essential role in GTPase catalysis, which is required for function and dynamic turnover of TSF protein filaments ^25^. On the other hand, G108S caused complex behaviours, where relatively large assemblies were stabilized in vivo, but smaller in vivo assemblies and in vitro polymers displayed reduced stability compared to wild-type CetZ1. This is consistent with at least two impacts of G108S on CetZ1 polymerization, reminiscent of the behaviour of FtsZ.G105S in *E. coli* ^13^, where GTPase activities are reduced thus stabilizing the GTP-hydrolysis mediated turnover of larger assemblies, and yet the longitudinal interface is also affected in a different manner, such that the protofilaments or other small assemblies are intrinsically less stable, independent of GTPase activity. Such behaviour suggests cooperative stabilization of large assemblies ^38^, which might involve other molecules in vivo that regulate or influence assembly, as seen with tubulin and FtsZ ^40,41^.

Related mutations in the GTPase active site of tubulin also reduce GTPase activity that alter microtubule polymerisation and depolymerisation ^42,43^. Our results with CetZ1 GTPase longitudinal interface mutations thus demonstrate the importance of these regions in CetZ1 polymerization and cellular function and confirm the evolutionary conservation of basic polymerization characteristics across the tubulin superfamily.

Our set of mutations targeting the predicted lateral association interface showed that this region is also important to CetZ1 polymerization and function overall. The most drastic effects were seen with a double mutation (R276E/K277E) that switches the charge distribution in a notably positively-charged region on the otherwise highly negative-charged CetZ1 molecule and involves residues expected to make direct contacts with the adjacent molecule in the sheet-like polymer model (Figure 2). Another more moderate substitution within the predicted lateral interaction interface (R282A) had a moderate effect on assembly and localization, but still strongly inhibited rod development. Other mutations in the vicinity of the lateral interaction site but not predicted to make direct lateral contact in the model (i.e., N269A and A295S) did not dramatically affect subcellular localization patterns but still partially inhibited rod development. These findings are consistent with an important role for lateral association in proper CetZ1 assembly and likely its cooperativity ^38^. However, the sensitivity of rod development function to some of the mutations in this region without drastic consequences to localization patterns might suggest that this region has additional functions affected by the mutations that are distinct from CetZ1’s function subcellular localization.

One region of interest outside the predicted longitudinal and lateral interfaces (Figure 2) was the unusually large Microtubule (M-) loop, a structural feature of TSF proteins named after its function in stabilizing lateral association on the interior surface of microtubules ^44^. Our results suggested that the conspicuous hydrophobic residues in an amphipathic region of CetZ1’s M-loop serve an essential function in CetZ1 polymerization, strongly impacting in vivo structures, dynamics, and envelope association, and showed congruent effects in vitro. This was despite the M-loop’s predicted significant protrusion from the surface of the globular TSF fold that mediates polymerization. We speculate that the long CetZ1 M-loop allows it to flexibly interact with adjacent subunits, analogous to the role of the different M-loop in tubulin ^45^ but likely via a differing structural mechanism due to its very different size and sequence; high-resolution structures of CetZ1 assemblies will be valuable in resolving its exact role in polymerization.

The E188G substitution on the ‘front’ surface (Figure 2) was important as it increased dynamic turnover of subcellular localizations and the frequency of small foci, and reduced polymer size in vitro, yet it showed full or slightly elevated function compared to wild-type in rod cell development. These findings point to a molecular mechanism where the dynamics of polymer behaviour are more instrumental in rod development than their static structures. Conversely, the mutants that caused stabilization of assembly dynamics strongly prevented rod development. This suggests the main role of CetZ1 is to dynamically guide a biosynthetic activity or the assembly of other rigid components at the cell envelope to reshape the cell, rather than to provide polymer rigidity directly. This model is also supported by our observation that CetZ1 wild-type polymer dynamics are much faster than the morphogenic transitions they influence. Such a mechanism is comparable to the proposed role for FtsZ during bacterial division constriction in guiding biosynthesis of the invaginating peptidoglycan cell wall ^46,47^.

The in vivo molecular and atomic-level structures of functional CetZ1 assemblies remain to be uncovered. Our findings suggest that CetZ1 does not form functional microtubules during rod development, but instead suggested that CetZ1 can polymerize cooperatively as bundled or sheet-like assemblies, which might directly associate with the membrane surface in *H. volcanii* ^18^. Previous work with overproduced CetZ1-GFP revealed infrequent very bright and dynamic cytoplasmic polymers that were highly reminiscent of microtubules and their classical dynamic instability behaviour ^18^, but are clearly different from the rod-development associated structures we have characterized here. We predict these are artefactual or rarely utilized microtubules occasionally formed during overproduction by tubulation of laterally-associated protofilaments in sheet-like assemblies, where based on our results, one might hypothesise the CetZ1 M-loop plays a role as it does in stabilizing classical tubulin microtubules ^12^.

The tubulin superfamily includes a wide range of proteins with distinct polymerization behaviours. The recently characterised Asgard archaeal OdinTubulin, for example, generates wider filament bundles compared to the canonical eukaryotic tubulins ^48^. The diversity in characterized polymerization behaviours, which now includes CetZs, highlights the adaptability of the TSF fold to multiple higher-order assembly modes that are essential to their roles. Our study also contributes to realizing the importance of dynamic filament turnover and cytomotive polymer behaviours that appear to drive the activities of many cytoskeletal proteins in the cellular context ^49^. Further comparative investigations of diverse cytoskeletal systems, including archaeal CetZ, FtsZ, and tubulin homologs ^8,50,51^, are expected to uncover common mechanisms and differences that could be exploited for engineering applications and should help resolve questions in cell biology including early evolutionary transitions in prokaryotic and eukaryotic cell form and function.

## Acknowledgements

This project was supported by the Australian Research Council (DP160101076 and FT160100010). We thank Mark Lockrey (TEM, UTS Microstructural Analysis Unit), Louise Cole and Amy Bottomly (UTS Microbial Imaging Facility), and Katharine Michie (UNSW, MWAC Structural Biology Facility) for technical and facilities management support, and Katharine Michie and Jan Löwe for scientific discussions. Dedicated to the memory of Linda Amos (1943-2021) and in acknowledgement of interesting discussions shared during the preliminary stages of this study.

## Author Contributions

Conceptualization – IGD, Data curation – RTdS, VS, HJB, IGD, Formal Analysis – RTdS, VS, HJB, Funding Acquisition – IGD, Investigation – RTdS, VS, HJB, IGD, Methodology - RTdS, VS, HJB, IGD, Supervision/mentoring – YL, IGD, Validation – VS, HJB, Visualization – RTdS, VS, HJB, IGD, Writing – RTdS, VS, HJB, IGD, Reviewing and editing – all authors. Estimates of overall author contributions (CalculAuthor weighted contributions method): RdS – 31%, VS – 23%, HJB – 21%, YL – 2%, IGD – 23%.

