## Supplementary Information for "Dynamic self-association of archaeal tubulin-like protein CetZ1 drives *Haloferax volcanii* morphogenesis"

### Contents

**Supplementary Figure 1.** Western blotting analysis of CetZ1 and CetZ1 mutant fusions with mTurquoise2.

**Supplementary Figure 2.** Focus thresholding and tracking and of CetZ1-mTq2.

**Supplementary Figure 3.** SDS-PAGE analysis of CetZ1 polymerization and lipid binding.

**Supplementary Video Legends**

**Supplementary Table 1.** Primers used in this study.

**Supplementary Table 2**. Plasmids used in this study.

**Methods for Supplementary data**

*Ultracentrifuge Polymer Pelleting Assay*

*Lipid Extraction and Purification from H. volcanii*

*Co-pelleting Assay for Lipid Binding*

**References**

### Supplementary Figures

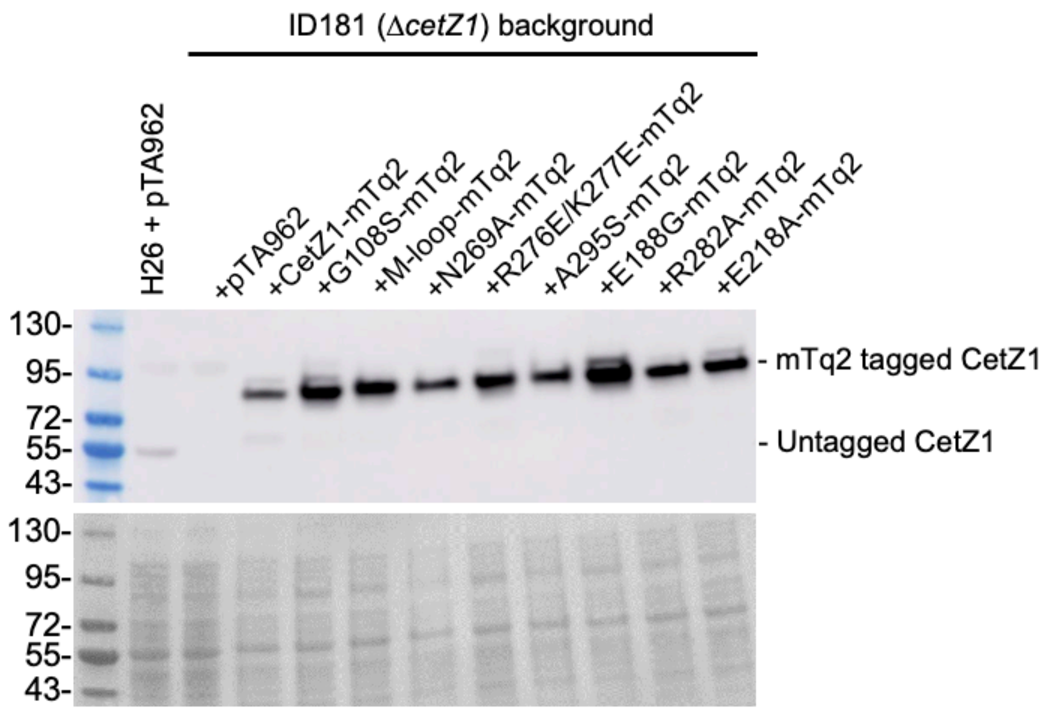


**Supplementary Figure 1. Western blotting analysis of CetZ1 and CetZ1 mutant fusions with mTurquoise2.** Strain ID181, the *cetZ1* knockout background, containing plasmids for the expression of mTq2 tagged CetZ1 and CetZ1 mutants was grown in Hv-Cab medium supplemented with 2 mM L-Tryptophan to induce expression. Whole cell-lysates were prepared at an approximate OD of 0.4 for SDS-PAGE and western blotting using an antibody against CetZ1 (top panel). H26 WT and ID181 containing the empty vector pTA962 were used as controls. Ponceau S staining was carried out before probing to ensure equal loading of samples before blotting (bottom panel).

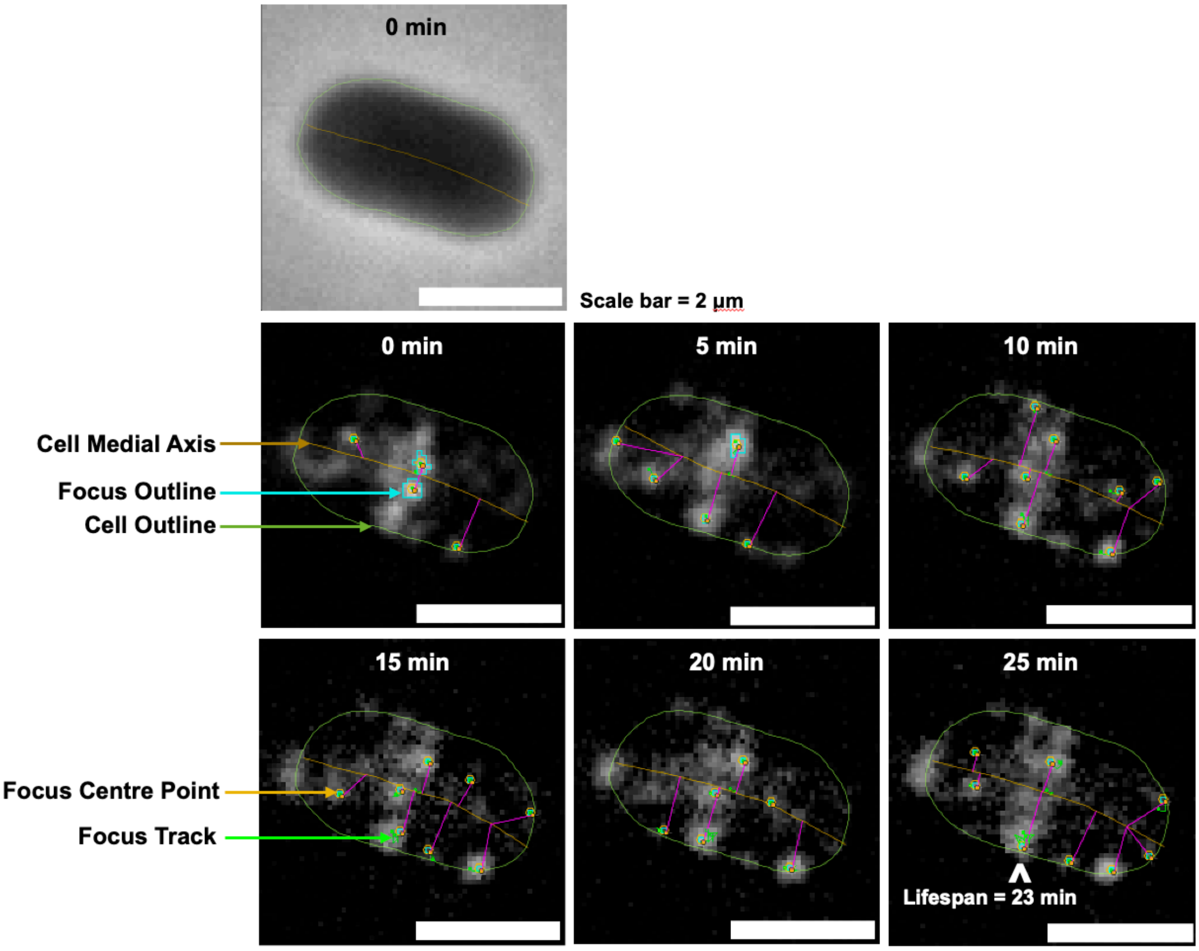


**Supplementary Figure 2. Focus thresholding and tracking and of CetZ1-mTq2.** Cell outlines (green) and the medial axis (brown) were determined using default thresholding of phase-contrast images (right) (see Methods). Thresholding of fluorescence images for foci detection, showing their outline (cyan), centre point (orange hexagon). Cumulative traces of the focus track over its lifespan are also shown for tracked foci (lime green). This example features a single cell of Δ*cetZ1* (ID181) + CetZ1-mTq2 from the data shown in Figure 5.

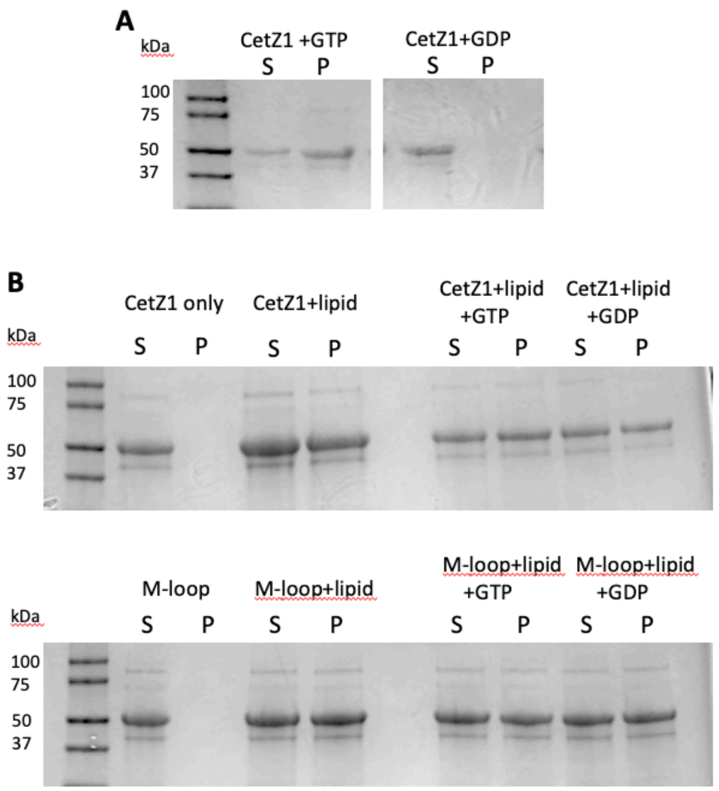


**Supplementary Figure 3. SDS-PAGE analysis of CetZ1 polymerization and lipid binding.** (A) CetZ1 (12 µM) reactions including 2 mM GTP or GDP in CetZ1 polymerization buffer were separated by ultracentrifugation into soluble (S) and pellet (P) fractions and analysed by SDS-PAGE. (B) Lipid association by CetZ1 and the M-loop mutant. CetZ1, with or without lipids and GTP/GDP in liposome buffer were mixed and subjected to ultracentrifugation to separate the supernatant (S) and pellet (P) fractions before SDS-PAGE. Preliminary experiments substituting CetZ1 with BSA revealed no protein in the pellet.

### Supplementary video legends

**Supplementary Video 1.** CetZ1-mTq2 localization throughout rod development via long-term time-lapse imaging (18 h, 10 min intervals) of CetZ1-G-mTq localization in cells transitioning from plate to rod shape.

**Supplementary Video 2.** 3D renders of CetZ1-mTq2 localization during mid-log phase. Scale bar: 0.5 µm.

**Supplementary Video 3**. 3D render of M-loop-mTq2 localization during mid-log phase. Scale bar: 0.5 µm.

**Supplementary Video 4.** Time-laps imaging of CetZ1-mTq2 during early-log phase growth in Hv-Cab medium supplemented with 2 mM L-Tryptophan. Images were taken every 1 min for 30 min. Scale bar: 5 µm.

**Supplementary Video 5.** Time-laps imaging of CetZ1.E218A-mTq2 during early-log phase growth in Hv-Cab medium supplemented with 2 mM L-Tryptophan. Images were taken every 1 min for 30 min. Scale bar: 5 µm.

**Supplementary Video 6.** Time-laps imaging of CetZ1.G108S-mTq2 during early-log phase growth in Hv-Cab medium supplemented with 2 mM L-Tryptophan. Images were taken every 1 min for 30 min. Scale bar: 5 µm.

**Supplementary Video 7.** Time-laps imaging of CetZ1.Mloop-mTq2 during early-log phase growth in Hv-Cab medium supplemented with 2 mM L-Tryptophan. Images were taken every 1 min for 30 min. Scale bar: 5 µm.

**Supplementary Video 8.** Time-laps imaging of CetZ1.E188G-mTq2 during early-log phase growth in Hv-Cab medium supplemented with 2 mM L-Tryptophan. Images were taken every 1 min for 30 min. Scale bar: 5 µm.

### Supplementary Tables

**Supplementary Table 1.** Primers used in this study

| Primer | Sequence (5'-3') |
|---|---|
| CetZ1_N269A_F | caccgcccacacgaccgcgcgaatcacgagcctc |
| CetZ1_N269A_R | gaggctcgtgattcgcgcggtcgtgtgggcggtg |
| CetZ1_R276E/K277E_F | gaatcacgagcctcgtcgaggaggccgcgctcggtc |
| CetZ1_R276E/K277E_R | gaccgagcgcggcctcctcgacgaggctcgtgattc |
| CetZ1_A295S_F | gaaggcgcggagcgctcgctgctcgtgctcgctg |
| CetZ1_A295S_R | cagcgagcacgagcagcgagcgctccgcgccttc |
| CetZ1_E188G_F | gatacgacgagatcaacggcgaaatcgtcaaccg |
| CetZ1_E188G_R | cggttgacgatttcgccgttgatctcgtcgtatc |
| CetZ1_R282A_F | aaggccgcgctcggtgcgctcacgctcccgtg |
| CetZ1_R282A_R | cacgggagcgtgagcgcaccgagcgcggcctt |
| CetZ1_G108S_F | cgtgtccgggctctcgggcggcaccggctc |
| CetZ1_G108S_R | gagccggtgccgcccgagagcccggacacg |
| CetZ1_M-loop HPM_F | ggaagaacaacggcggcggctcggcgtctcgggcgacgggcggcgacgagc |
| CetZ1_M-loop HPM_R | gctcgtcgccgcccgtcgcccgagacgccgagccgccgccgttgttcttcc |
| CetZ1_C-term tailM_R (stop) | cgcggatccttagccgcccgactccgcttcgtcctcgtcatcgtttatgagg |
| CetZ1_C-term tailM2_R (no stop) | cgcggatccgccgcccgactccgcttcgtcctcgtcatcgtttatgagg |
| EndP_CetZ1_F | cccccgggaattcatatgaagctcgcaatgatcggattcggg |
| EndP_CetZ1_R (stop) | cgcggatccttagaaaagcgactccagttcgtcctcg |
| EndP_CetZ1_2_R (no stop) | cgcggatccgaaaagcgactccagttcgtcctcg |

**Supplementary Table 2**. Plasmids used in this study

| Plasmid | Description | Source |
|---|---|---|
| pTA962 | *H. volcanii – E. coli* shuttle expression vector | [1] |
| pTA962-CetZ1 | For expression of untagged CetZ1 | [2] |
| pTA962-CetZ1.E218A | For expression of untagged CetZ1.E218A | [2] |
| pTA962-G108S | For expression of untagged CetZ1.G108S | This study |
| pTA962-E188G | For expression of untagged CetZ1.E188G | This study |
| pTA962-Mloop | For expression of untagged CetZ1.Mloop | This study |
| pTA962-CTT | For expression of untagged CetZ1.CTT (C-terminal tail) | This study |
| pTA962-R282A | For expression of untagged CetZ1.R282A | This study |
| pTA962-N269A | For expression of untagged CetZ1.N269A | This study |
| pTA962-R276E/K277E | For expression of untagged CetZ1.R276E/K277E double mutant | This study |
| pTA962-A295S | For expression of untagged CetZ1.A295S | This study |
| pHVID9 | Vector backbone used to generate G-linker mTurquoise2 C-terminal fusion proteins | [3] |
| pHVID135 | For expression of CetZ1-mTq2 C-terminal fusion with G-linker | [3] |
| pHJB13 | For expression of CetZ1.E218A-mTq2 C-terminal fusion with G-linker | This study |
| pHVID9-G108S | For expression of CetZ1.G108S-mTq2 C-terminal fusion with G-linker | This study |
| pHVID9-E188G | For expression of CetZ1.E188G-mTq2 C-terminal fusion with G-linker | This study |
| pHVID9-Mloop | For expression of CetZ1.Mloop-mTq2 C-terminal fusion with G-linker | This study |
| pHVID9-R282A | For expression of CetZ1.R282A-mTq2 C-terminal fusion with G-linker | This study |
| pHVID9-N269A | For expression of CetZ1.N269A-mTq2 C-terminal fusion with G-linker | This study |
| pHVID9-R276E/K277E | For expression of CetZ1.R276E/K277E-mTq2 C-terminal fusion with G-linker | This study |
| pHVID9-A295S | For expression of CetZ1.A295S-mTq2 C-terminal fusion with G-linker | This study |

| | | |
|---|---|---|
| pHIS17 | Vector backbone used to generate plasmids for expression and purification of CetZ1 and its mutants in *E. coli* | [4] |
| pHIS17-CetZ1 | For production and purification of CetZ1 from *E. coli* | This study |
| pHIS17-E218A | For production and purification of CetZ1.E218A from *E. coli* | This study |
| pHIS17-G108S | For production and purification of CetZ1.G108S from *E. coli* | This study |
| pHIS17-E188G | For production and purification of CetZ1.E188G from *E. coli* | This study |
| pHIS17-Mloop | For production and purification of CetZ1.Mloop from *E. coli* | This study |
| pHIS17- R276E/K277E | For production and purification of CetZ1 R276E K277E double mutant from *E. coli* | This study |

**Methods for Supplementary Data**

***Ultracentrifuge Polymer Pelleting Assay***

A reaction mixture was prepared with 12 µM CetZ1 or mutant in CetZ1 polymerization buffer (50 µL). The samples in micro-ultracentrifuge tubes were incubated for 2 min with shaking at 300 rpm at 37 °C. Polymerization was initiated by adding 1 µL of 100 mM GTP and further incubated for 2 min. Subsequently, the tubes were transferred to a Beckman TLA100 rotor and centrifuged for 10 min at 90,000 rpm and 20 °C. The supernatant was carefully transferred into a clean tube, and the pellet was gently washed with CetZ1 polymerization buffer, solubilized with SDS sample buffer, and analyzed by SDS-PAGE.

***Lipid Extraction and Purification from H. volcanii***

The method was based on those previously described [5,6]. *H. volcanii* cells were harvested during the late exponential growth phase by centrifugation at 7,000 rpm for 10 min. The cells were washed by re-suspending in 50 mL of 18 % BSW and centrifuged at 6,500 rpm for 10 min. The supernatant was discarded, and the pellet was washed with 50 mL of 18 % BSW and subsequently freeze dried (24 h). To extract lipids the freeze-dried cells were resuspended in a mixture of methanol, dichloromethane, and phosphate buffer (8.7 g/L $KH_2PO_4$, pH 7.4) at a ratio of 2:1:0.8 (v/v) (5 mL per gram), then sonicated in a water bath sonicator (Power Sonic 420, 40 kHz, 700 W) for 5 min. The sample was centrifuged at 6,500 rpm for 5 min and the supernatant was isolated. The extraction procedure was repeated on the pellet and the supernatant was pooled with the first extract. Two additional extractions were then performed on the remaining pellet, using 0.8 volumes of 0.5 % Trichloroacetic acid (pH 2).

To further purify lipids, 0.5 volumes of dichloromethane and ultrapure water were added to the pooled extracts, vortexed and centrifuged at 6,500 rpm for 5 min. After the phases had visibly separated, the organic phase was retained, and the aqueous phase was then washed by adding dichloromethane (5 mL per gram of original dry mass) followed by vortexing and centrifugation. This second dichloromethane phase was combined with the main organic phase. Finally, the organic phase was washed with equal volume of pure water, centrifuged and the organic phase was collected. The final organic phase was aliquoted in glass vials and evaporated under a stream of dry $N_2$ gas. The vials were sealed while under $N_2$ flow to minimize the oxygen in the vial and stored at -20 °C.

***Co-pelleting Assay for Lipid Binding***

The liposomes were prepared by re-suspending the extracted lipids in filter sterilized liposome buffer (50 mM Tris-Cl, 200 mM KCl, 1 mM EDTA pH 8.5), adding 5 mL of liposome buffer per 10 mg of lipids. The mixture was vortexed for several seconds, then sonicated in a water bath sonicator (40 kHz, 700 W) for 30 min. Liposomes were stored at 4 °C and used within 7 days.

The co-pelleting assay was performed using sonicated liposomes. Purified CetZ1 was first centrifuged using Beckman TLA 100 rotor at 50,000 rpm to remove any aggregated protein. Liposomes (44 µL at 2 mg/mL) were mixed with 5 µL pre-spun CetZ1 (to give final CetZ1 concentration of 12 µM) in liposome buffer, and, where indicated, 1 µL of 100 mM GTP/GDP was mixed in and incubation was continued for 2 min. The mixture was then centrifuged at 55,000 rpm at 20 °C for 25 min. The supernatant and pellet were isolated for analysis by SDS-PAGE.
